# Selecting the most important self-assessed features for predicting conversion to Mild Cognitive Impairment with Random Forest and Permutation-based methods

**DOI:** 10.1101/785519

**Authors:** Jaime Gómez-Ramírez, Marina Ávila-Villanueva, Miguel Ángel Fernández-Blázquez

## Abstract

Alzheimer’s Disease (AD) is a complex, multifactorial and comorbid condition. The asymptomatic behavior in the early stages makes the identification of the disease onset particularly challenging. Mild cognitive impairment (MCI) is an intermediary stage between the expected decline of normal aging and the pathological decline associated with dementia. The identification of risk factors for MCI is thus sorely needed. Self-reported personal information such as age, education, income level, sleep, diet, physical exercise, etc. are called to play a key role not only in the early identification of MCI but also in the design of personalized interventions and the promotion of patients empowerment. In this study we leverage on *The Vallecas Project*, a large longitudinal study on healthy aging in Spain, to identify the most important self-reported features for future conversion to MCI. Using machine learning (random forest) and permutation-based methods we select the set of most important self-reported variables for MCI conversion which includes among others, subjective cognitive decline, educational level, working experience, social life, and diet. Subjective cognitive decline stands as the most important feature for future conversion to MCI across different feature selection techniques.

## Introduction

Mild cognitive impairment (MCI) is an intermediary stage between the expected decline of normal aging and the pathological decline associated with dementia. MCI patients compared with cognitively normal cases have a higher risk of progressing to dementia^1,2,3,4,5^. Although the diagnosis of MCI has prognostic value, the value is intrinsically linked to the time of the diagnosis. The early detection of MCI is thus a priority target in aging research^6^.

There is not only a concern with the early detection of cognitive decline symptoms, it is also important to understand the time stability of the MCI diagnosisc,^7,8^. The *yo-yo effect* refers to fluctuations between normal and MCI diagnoses observed for the same person. The variability in symptoms may lead to spurious diagnosis, for instance, a neurologist or a neuropsychologist may diagnose a person with MCI and later in the future retract the diagnosis9. Longitudinal studies are best equipped to deal with *yo-yo effects* and gain an overall view of personal trajectories of disease progression.

To better understand this variability, we need to take into account several factors that may have a measurable effect on cognitive performance. MCI is, in the end, a clinical description based on performance on tests of memory and thinking skills. The capacity to perform well on a test could be affected by lifestyle conditions. For example, a person going through stressful events, sleep-deprived or with a poor diet may have worsening memory test scores compared to previous occasions with more favorable personal circumstances^10,11,12,13^.

Testing cognition in a large elderly population on a regular basis, rather than when memory loss starts to occur, might help us understand the role played by fluctuations between normal and MCI conditions in the risk of later developing dementia. The systematic examination via, for example, cognitive testing, brain imaging techniques or gene expression profiling, of a very large of pool subjects, is in all cases extremely costly. Prior or in addition to such an effort it is worth collecting information variables that can be directly assessed by the individuals and demonstrably have an effect on cognitive impairment, such as lifestyle, diet, sleep patterns or subjective memory decline.

In this work, we use the dataset collected in *The Vallecas Project*, an ambitious longitudinal community-based study for healthy aging in Spain. For the reasons above discussed, we focus on features that can be self-assessed by the participants. Self-assessed features are variables that can be reported by the subject herself (e.g. age, income level, education, sleep, diet, physical exercise, etc.). Our goal is to study the most important features to predict conversion to MCI using Random Forest and permutation-based techniques that help us understand the real effect of the predictors (self-reported variables) in the target (conversion to MCI).

## Methods

*The Vallecas Project* is an ongoing single-center, observational, longitudinal cohort study^14,15^. The participants, recruited between 2011 and 2013 in Madrid, Spain, are home-dwelling volunteers, aged 70 to 85, without relevant psychiatric, neurological or systemic disorders. After signing informed consent, they undertake a yearly systematic clinical assessment including medical history, neurological and neuropsychological exam, blood collection and brain MRI. The cohort size was 1,180 subjects at baseline. Since then, the number of active subjects has decreased across the years, 964 subjects came to the second visit, 865 the third visit, 773 the fourth visit, 704 the fifth visit and 509 to the sixth visit, the last yearly visited completed. At the time of this writing (07/09/2019) the project is running the 7th and 8th visits.

The main objective of *The Vallecas Project* is to elucidate the best combination of features that are informative about developing cognitive impairment in the future. The subjects in each visit were diagnosed as healthy, mild cognitive impairment (MCI) or dementia. *The Vallecas Project* dataset includes information about a wide range of factors including magnetic resonance imaging (MRI), genetic, demographic, socioeconomic, cognitive performance, subjective cognitive decline, neuropsychiatric disorders, cardiovascular, sleep, diet, physical exercise and self-assessed quality of life.

In this work, we focus on features that are self-assessed by the participants in *The Vallecas Project* for the completed visits, that is, from visit first to sixth. Specifically, the features of interest fall within the following categories: demographics, anthropometric, neuropsychiatric, traumatic brain injury, cardiovascular, quality of life, engagement with the external world, physical exercise, social engagement, sleep, diet, and subjective cognitive decline. Table 1 shows the types of self-assessed features collected in *The Vallecas Project* and studied here. A complete description of the dataset is included in the Supplementary Materials, Table S1.

**Table 1.** Feature types, each category contains several features.

|  |  |
| --- | --- |
| Demographics | Anthropometric |
| Neuropsychiatric | Diet |
| Cardiovascular | Quality of Life |
| Engagement External World | Physical Exercise |
| Social Engagement | Traumatic Brain Injury |
| Sleep | Subjective Cognitive Decline |

### Automated Feature Selection

We are interested in studying the predictive power of self-assessed features collected in *The Vallecas Project* on future conversion to mild cognitive impairment (MCI). This is a feature selection problem aiming at detecting the most important features to predict conversion to MCI. The engine of the automatic feature selection problem has as input the set of self-assessed features (Table S1) measured in year 1 and as output target, the conversion to MCI diagnosis in the latest available visit (year 2 to year 6 both inclusive).

If properly tackled, the problem of feature selection needs to deal with at least three milestones. First, *How many* features must be included in the minimum set of important features; second, *Which* are the most important features and third; *Why* are those features the most important ones. Thus, we must address how many features we need to consider, identify which are those and finally explain why those features are important in terms of prediction.

#### How many features? The one in ten rule

The first question to be pondered is, How much data is enough to consistently predict the target? The answer is not straightforward and depends on a number of factors, for example, the type of model (linear or non-linear), the accuracy we want to achieve, the quality of the data (signal-to-noise ratio), the number of inputs and so on. The required size of the training data is thus an ill-posed question, however, heuristics that address this problem are available. One such heuristic is the *one in ten* rule which states that the amount of data needed is 10 times the number of parameters in the model16. For example, according to the *one in ten* rule of thumb, in a sample of 1,000 subjects with 140 positive cases (e.g converted to MCI), 14 parameters, that is, the 10% or 1 in 10 of the minority class, can be used to reliably fit the total dataset.

The *one in ten* rule effectively transforms the problem of deciding the size of the training set by that of knowing the number of parameters in the model. In the case of linear models, this is trivial since the number of parameters is equal to the number of inputs. Nevertheless, the *one in ten* rule should be seen as a reasonable guess on the number of features and never as a prerequisite.

#### Which features?

As important or more as deciding about the number of features to be included in the model is to be able to asses the relative importance of the features. In order to study which are the most important features for prediction accuracy, we need to discuss first the required methodology to estimate the usefulness of the features. Depending on the evaluation metric, we can distinguish between two methodologies for automated feature selection: *filter* and *embedded methods*. Both methodologies are apt to be used to remove the non-essential features for the task of predicting new values of the target feature, that is, the *Which are the important features* question. Filter methods and embedded methods (random forests) are introduced next.

Filter methods pick up the intrinsic properties of the features estimated via univariate correlation matrix. In essence, a filter method is a linear approach to finding variables that contain information about the target variable by means of statistical tests (e.g. chi-squared test, Fisher’s exact test) or related quantities such as the correlation coefficient.

The algorithm *SelectKBest*^17^ *is one possible implementation of the filter method. SelectKBest* uses a score function in order to remove all but the highest scoring features, that is to say, only the *k* most important features are retained. The algorithm can take as score function the Fisher’s test, the 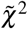 and the mutual information. For example, the referred algorithm with 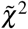 as the score function selects the *k* most important input features *X* to predict the target feature *y*. A small value of the 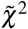 statistic for (*x* ∈ *X, y*) means that the feature *x* is independent of *y*. On the other hand, a large value means that *x* is non-randomly related to *y*, and therefore likely to contain information about target *y*.

Random forest is one of the most popular machine learning algorithms. Random forests tend to perform well in small and medium-size datasets, and most importantly for this study, random decision forest is the algorithm of choice for automated feature selection^18^. Random forests solve the real problem of feature selection because it tells us which are the most important features to optimize the prediction. Filter methods, on the other hand, do not solve any optimization problem, they rather tell us which features are most linearly correlated with the target feature.

How ensemble methods such as random forest work may be understood by using the analogy of asking a thousand people (experts) and then aggregating their answers. Likely, the aggregated response is better than the individual expert’s responses. By the same token, the aggregation of many inaccurate predictors (forest) will give better answers than the individual predictions (tree). Or going from analogy to allegory: *Don’t look at the tree, look at the forest instead!*

A random forest consists of a large number of decision trees (tree-like model of decisions) where each tree is trained on *bagged data* (sampling with replacement) using random selection of features19. Thus, a random forest is, in essence, a meta estimator that fits a number of decision tree classifiers. Decision trees are nonparametric models, that is to say, the model will have as many parameters as it needs to fit the data. It follows that if left unconstrained, the tree will fit the data very closely, most likely overfitting. Decision trees are unstable in the sense that small changes in the input may produce very different decision trees.

Although in principle, random forests do not suffer from multi-collinearity issues due to highly correlated features, it is, however, advisable to take care of redundancy before training a random forest. As a matter of fact, having a large set of variables containing similar information may induce the model to weigh heavily on this set in detriment of others^20^. Random forests effectively address the overfitting and the stability problems existing in decision trees. Furthermore, random forests do not suffer from the limitations of the beforementioned filter methods. Filter methods use correlation to assess the relevance of features, but they are likely to fail to find the best subset of features when features do not behave linearly, e.g. non-normality, multicollinearity or heterocedasticity^21^ exist in the data set.

The Gini importance -a computationally efficient approximation to information entropy^22^- is a score that provides a relative ranking of the spectral features and is a by-product of the training process in a random forest classifier, that is to say, the feature selection mechanism is embedded in the training algorithm of the classifier. To understand the Gini importance is necessary to understand first Gini impurity which is a measure used in decision trees to determine how often something is incorrectly labeled if that labeling were random. For example, if half of the data points are in class “A” and the other half in class “B”, a data point randomly chosen will have a 50% chance of being labeled incorrectly. Formally, the Gini impurity for a set of M classes with *p*_*i*_ the fraction of items labeled with class *i* shown in Equation 1. The Gini impurity reaches its minimum (zero) when all points fall into the same category.

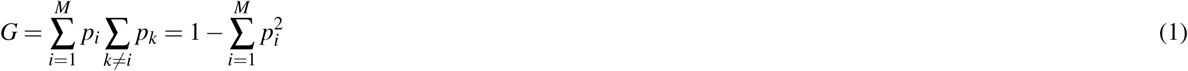

Now, since Random forest are an ensemble method of individual decision trees, the Gini impurity can be used to calculate the mean decrease in Gini across all trees or Gini importance. The Gini importance for a node is the average decrease in node impurity, weighted by the proportion of samples reaching that node in each individual decision tree in the random forest. A higher Mean Decrease in Gini indicates higher variable importance. The importance of node j, *I*(*j*), assuming only two child nodes (binary tree) is then calculated as:

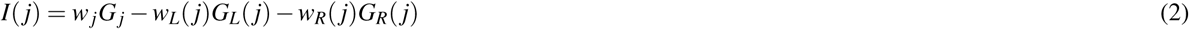

where *w*_*j*_*G*_*j*_ is the impurity value of node *j* weighted by number of samples reaching node *j, w*_*L*(*j*)_*G*_*L*(*j*)_ is the weighted impurity value of the child node from the left split on node *j* and *w*_*R*(*j*)_*G*_*R*(*j*)_ is the weighted impurity value of child node from the right split on node *j*.

#### Why the important features are important?

Once the *How many* and the *Which ones* questions have been addressed, there is one question left: *Why the important features are such?.* As it was shown in the Section above, random forests compute the importance of features as the mean decrease of the Gini impurity. However, from a more fundamental standpoint, the importance of a feature can be seen as the increase in the prediction error of the model after we permute the feature’s values. By virtue of permuting the feature’s values, the relationship between the feature and the true outcome is broken. If the feature were important for model accuracy, the accuracy will worsen upon the permutation of the feature values. By the same token, is a feature is unimportant, shuffling its values will likely leave the model error unchanged.

The permutation feature importance measurement was introduced by Breiman^18,23^ for random forests, however, the procedure is model-agnostic and can be used for any other machine learning model. Feature importance can be assessed via permutation methods which in essence quantify the effect on model accuracy of randomly reshuffling each predictor variable. Permutation-based importance methods are a reliable technique that do not suffer from the bias existing in Gini impurity which might inflate the importance of continuous and high-cardinality categorical variables.

This approach directly measures feature importance by observing how random re-shuffling (thus preserving the distribution of the variable) of each predictor influences model performance. However, removing each feature from the dataset to then re-train the estimator is computationally very intensive. A more efficient approach that avoids retraining as many estimators as features was proposed by Fisher and colleagues^24^. The algorithm calculates the importance of features based on changes in the prediction error and is described below.

Input: Trained model F, Input feature matrix X, Target vector y and Error measure L.

Step 1: Estimate the original model error *ε*^*o*^ = *L*(*y, F*(*X*))
Step 2: For each feature *j* do:
  Step 2.a: Shuffle the values in *X*_*j*_ to obtain *X* ^*perm*^ (the original association between *X*_*j*_ and the target *y* is broken).
  Step 2.b: Estimate the new error based on the predictions of the shuffled data, *ε* ^*perm*^ = *L*(*y, F*(*X* ^*perm*^)).
  Step 2.c: Calculate the permutation feature importance of feature j (*I* ^*j*^) as the difference between the error before and after shuffling the values, *I* ^*j*^ = *ε* ^*perm*^ *-ε*^*o*^.
  Step 3: Sort the features, argmax _*j*_ *I*^*j*^.

The above algorithm measures the importance of features via the change in the model’s prediction error after permuting each feature. However, permutation feature importance does not contain information about how changes in the range of values of the variable change prediction.

The last permutation-based interpretation method discussed is the Shapley value method^25^. The Shapley value is the average marginal contribution of a feature value across all possible coalitions of feature values. For example, let us assume that we have trained a machine learning algorithm to predict conversion to MCI with three input features: Age, APOE and Income. A new patient comes in and the algorithm predicts that it will convert with probability 0.7, the average probability of conversion for all previous subjects is 0.3. We are asked to explain this prediction. To that end, we need to assess how much has each feature value contributed to the prediction compared to the average prediction. Linear regression models can address this question in a very straightforward way: the individual effects of each feature can be computed as the difference between the feature effect minus the average effect. However, if the assumption of linearity does not hold, which is the most likely scenario in complex real-world data, a different approach is needed.

The Shapley value method was originally developed in cooperative game theory^26^, and is apt for computing feature contributions for single predictions independently of the machine learning model used to fit the data. The Shapley value permits to calculate the contribution of a feature value as its contribution to the payout of the game of predicting the right label weighted and summed over all possible feature value combinations.

The Shapley value Φ is defined via a value function *ν* of all features in a set S. Specifically, the Shapley value of a feature value is its contribution to the payout (e.g. if the average prediction for all instances is 0.9 and the actual prediction is 0.8, the payout of 0.1) weighted and summed over all possible value combinations.

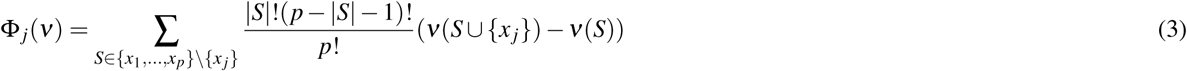

where *p* is the number of features, *S* is the subset of features, *x* is the vector of feature values of the particular instance to be explained and *ν*(*S*) is the prediction for feature values in *S*, marginalized over features that are not included in the set *S*.

The Shapley value is arguably the best permutation-based method for explaining the effects of feature values in the average prediction. Furthermore, Shapley value satisfies the properties of *Efficiency* (feature contributions must add up to the difference of prediction for an instance and the average), *Symmetry* (the contributions of two feature values are the same if they contribute equally to all possible coalitions), *Dummy* (a feature that does not change the predicted value has a Shapley value of 0) and *Additivity* (in the case of a random forest, for a feature value, the average of the Shapley value for each tree individually is equal to the Shapley value for the feature value for the random forest).

An example of the computation of Shapley Values is included in the Supplementary Material, also for a more in depth description of Shapley value, see^27^.

## Results

We are interested in selecting the most important self-assessed features for conversion to MCI for subjects that have at least 2 visits (920 subjects). The number of subjects that converted to MCI is 112 subjects (∼ 12.17% conversion rate (∼ 12.64% male, ∼11.89% female). The gender ratio in the study was 340 (37%) male and 580 (63%) female, the average age at the basal visit of the participants was 74.6±3.84, the body mass index 27.29±3.58 and the number of years of schooling of the participants 10.9±5.8 (12.42±6.12 male and 10.05±5.42 female). The total number of features considered to study conversion to MCI was 91 (Supplementary Materials Table S1). A complete description of the dataset is provided in^15^.

The automated feature selection problem is analyzed using the methods defined in the previous section-filter method, random forest feature importance, and permutation-based importance.

### Filter method

According to the *one in ten* rule discussed in Section, the number of features to be retained is 12 (10% of the number of cases in the positive class, 112). Figure 1 shows the 12 most important features based on the filter method *SelectKBest*^17^ which were in this order: APOE (APOE is not a self-assessed feature but it provides valuable information and allows us to have an idea about the importance of self-assessed variables compared to genetic variables), subjective cognitive decline (SCD), age, diet high in sweets, frequency family relationships, body mass index (BMI), self perceived deterioration in executive function, weight, thyroid-related problems, participation in cultural/art activities, difficulties remembering facts, and history of heart health (no heart problems, angina, infarct).

**Figure 1.**
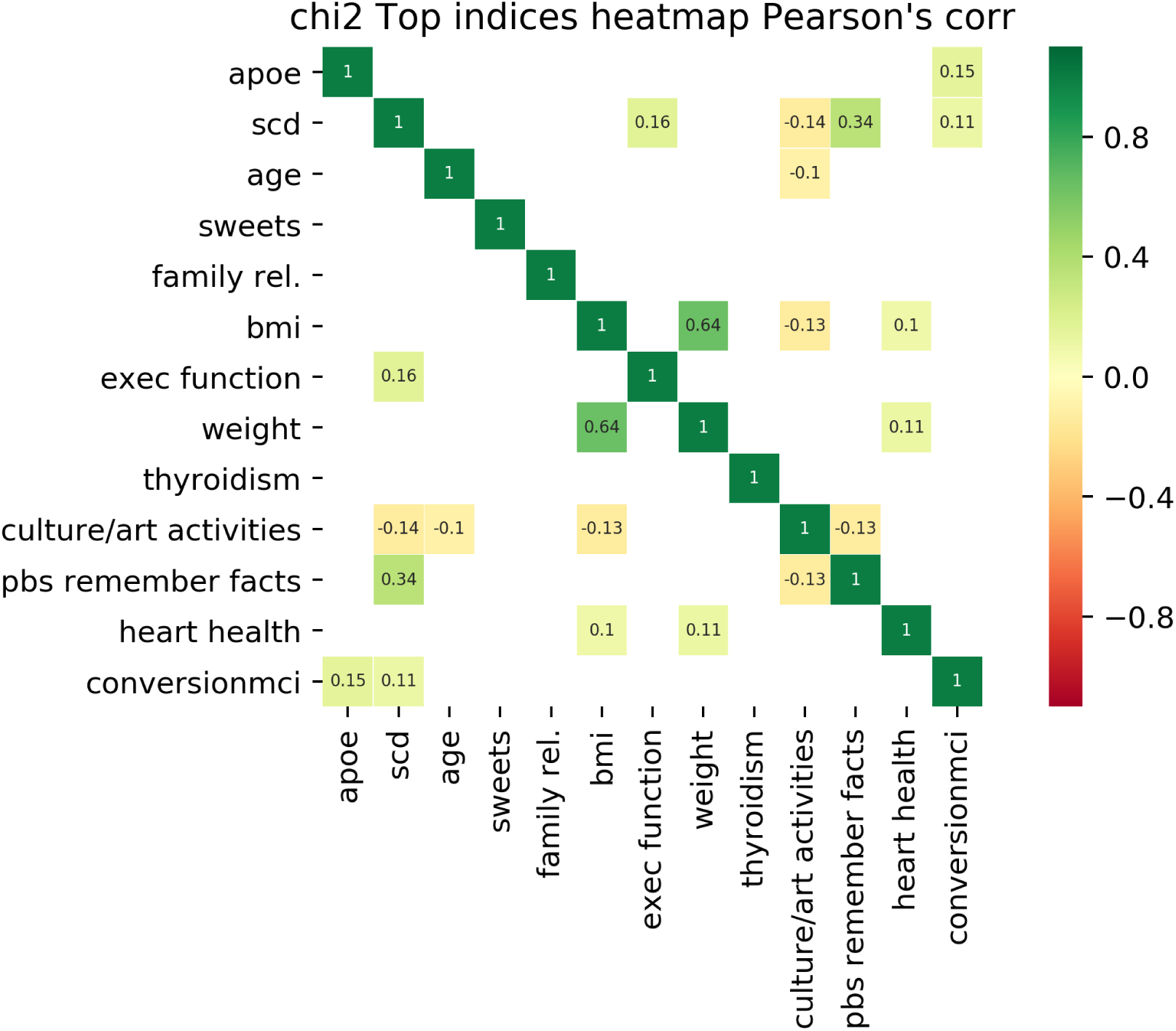
Evaluation of the importance of features for conversion to MCI classification using the filter method *SelectKBest*. The 12 features most informative for MCI conversion are displayed in order of importance (increasing importance from bottom to top or from left to right). The target feature, conversion to MCI, is displayed in the last row/column. Pearson’s correlation among all pairs of features including the output target (conversion to MCI) are shown when *r* > 0:1. Only APOE and SCD correlate above this threshold.

The Pearson’s correlation between features is also shown in the figure when *r* > 0.1. The largest Pearson’s correlation with conversion to MCI are APOE and subjective cognitive decline (SCD) *r*(*APOE, target*) = 0.15, *r*(*SCD, target*) = 0.11.

### Embedded method: Random forest

Embedded methods such as random forest have the evaluation metric built in the model during the learning process, in our case we build random forest that tries to maximize the accuracy score. Dummy classifiers provide a null metric and work as a sanity check on the model’s performance. Figure 2 shows the random forest accuracy and how it compares with three dummy predictors: random predictor, majority predictor, and minority predictor. The random predictors predict with equal probability that a subject has or has not MCI. The majority of predictors predicts always the majority class (not MCI), and conversely, the minority predictor always predicts the minority class (MCI).

**Figure 2.**
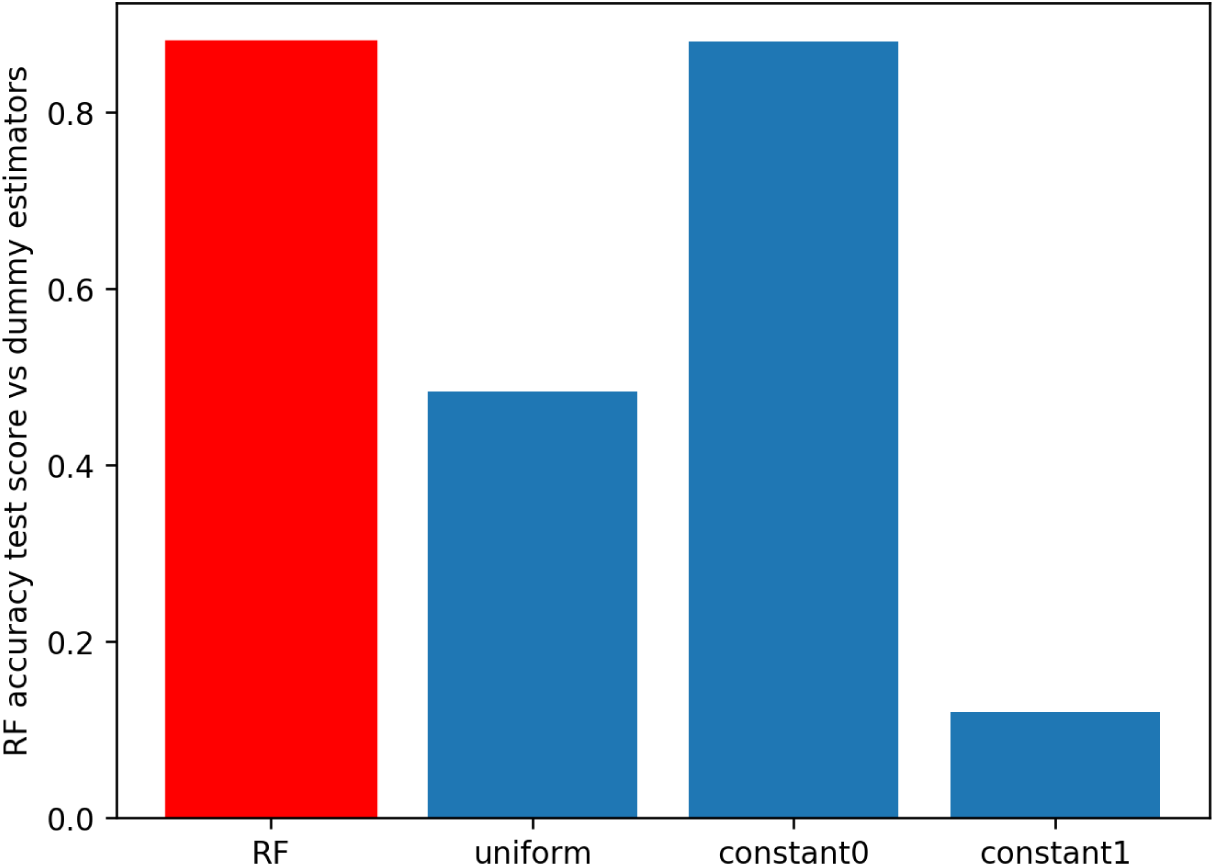
Comparative analysis of the accuracy score for the random forest predictor (in red) and 3 dummy predictors (in blue). The random forest shows a better accuracy score than the random classifier (uniform predictions) and the classifier that always predicts the minority class (constant1). The accuracy score of the random forest is 0.851, for the uniform predictor is 0.542, the majority predictor is 0.877 and for the minority predictor is 0.123.

The most important features as determined by the random forest are shown in Figure 3. Since the approaches for feature selection are different-linear vs. non-linear-it was expected that the set of important features varies from one method to the other. However, there was subset of features that was retained as the 12 most important in both methods, namely, subjective cognitive decline (SCD), years of schooling, body mass index (BMI), diet rich on sweets, age, relationship with friends and APOE. The subjective cognitive decline (SCD) was the most important feature in both methods.

**Figure 3.**
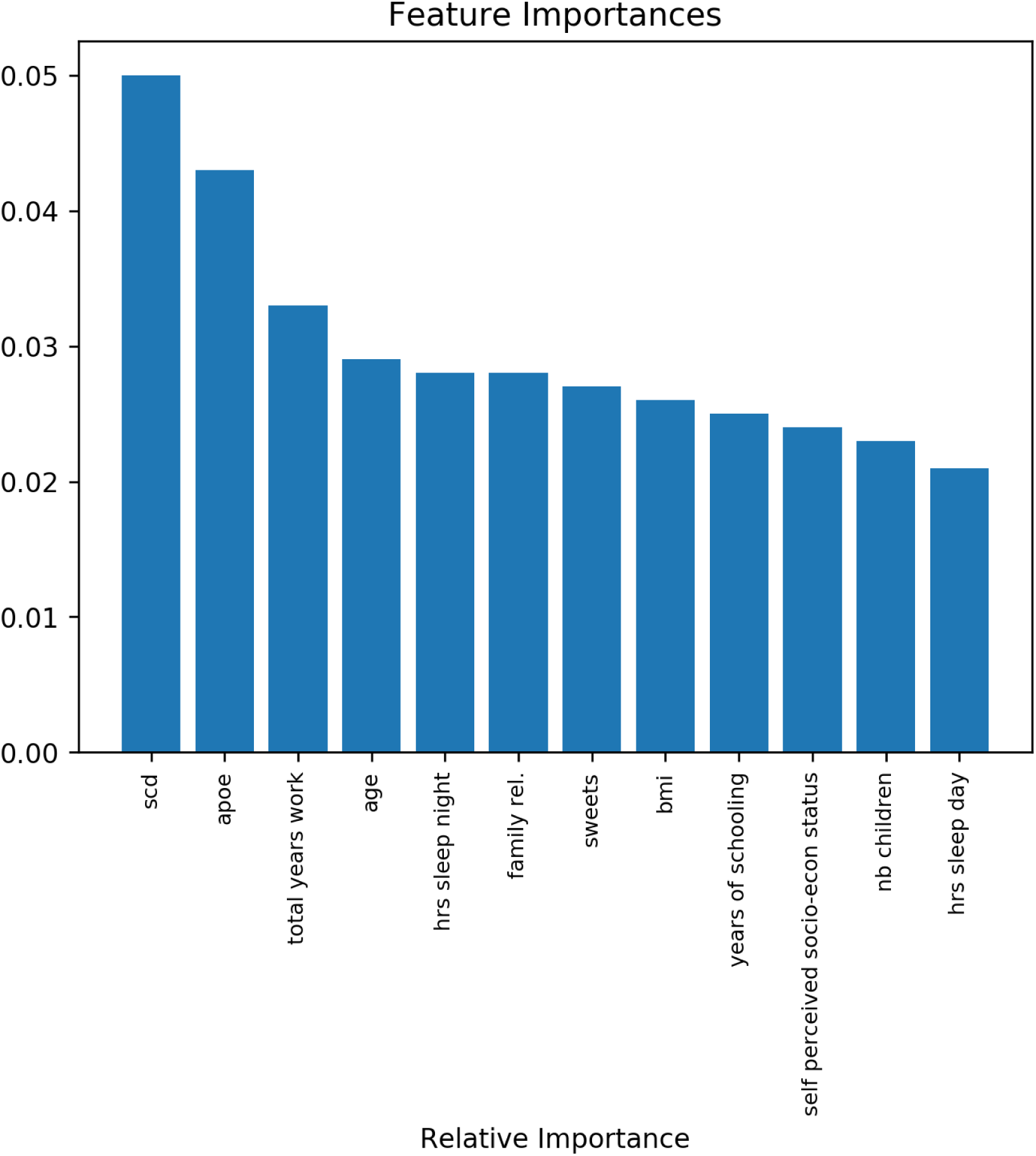
Evaluation of the importance of features for conversion to MCI. The 12 features most informative are displayed in the x-axis, the y-axis shows the importance of each feature based on the Gini impurity described in Equation 2. The sum of the Gini importance of all the features in the model is equal to 1, the figure shows only the top 12 features.

### Permutation-based importance methods

Figure 4 shows the output of the permutation importance algorithm which measures how the importance score decreases when a variable is shuffled and so breaking any prior relationship between variable and target. The features colored in green indicate that, as expected, the predictions of the shuffle data are less accurate than the real data. The red colored, on the other hand, indicate that the predictions of the shuffle data happened to be more accurate than the real data. Although this may seem surprising (by introducing noise we get better predictions), the rationale is uncomplicated, random chance caused the predictions of the noisy data to be more accurate than the actual values because the features most likely do not contain information about the target feature and the improve in the accuracy is purely coincidental and due to chance. The most important feature again is subjective cognitive decline (SCD).

**Figure 4.**
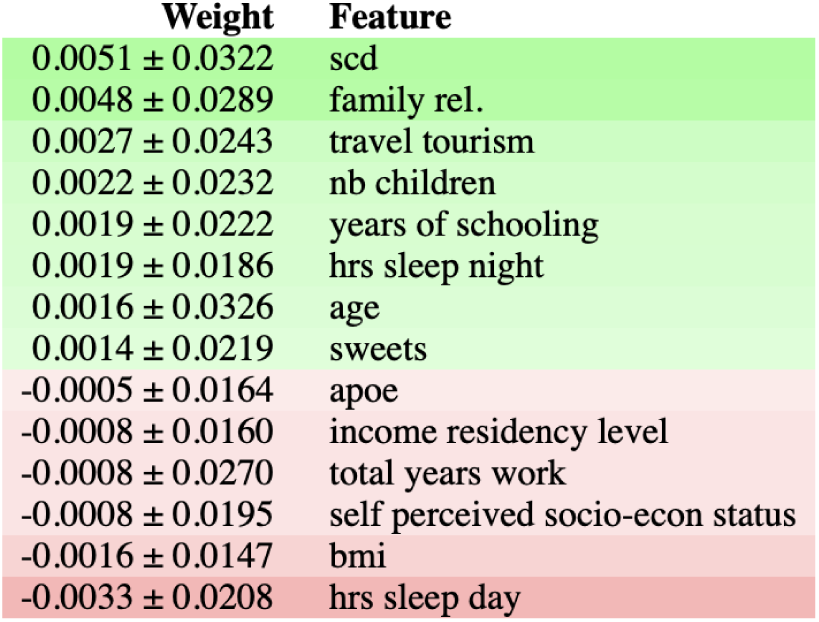
Figure shows feature importance using the ELI5 library^28^. The most important features are at the top and those at the bottom matter the least. The first column in each row shows how much model accuracy decreased with a random shuffling ± how the accuracy varied from one-reshuffling to the next. The most important feature is subjective cognitive decline (SCD), followed by diet features(sweets and white fish), hours of sleep during the day and APOE. The rows in red show predictions of the shuffle data that happened to be more accurate than the real data. The idea behind this is that random chance caused the predictions of the noisy data be more accurate, this indicates that the features do not contain information about the target feature.

Once we have studied the most important features according to the Gini score and analyzed the effect of random shuffling, it is possible to go a step further and look at the specific effect in prediction within the range of values of each feature. Partial Dependence Plots (PDP) separate out the effect of each feature on prediction. Figure 5 shows the PDP of four important features: SCD, APOE and Dietary Habits, sweets and white fish. The X-axis represents the range of values of the feature and the Y-axis shows changes in the prediction, positive values represent the contribution of the feature to the increase in the prediction (increase in the odds to convert to MCI) and negative values represent the contribution of the feature to the decrease the prediction (reduction in the odds to convert to MCI). For example, if *PDP*(*X* = 2) = 0.1 the feature X with value 2 increases 10% the chances to convert to to MCI, by the same token, if *PDP*(*x* = 2) = −0.1 the feature X with value 2 reduces the chances to convert to MCI 10%. The shaded area represent the standard deviation which is always 0 at the baseline point in the origin (*x* = 0, *y* = 0).

**Figure 5.**
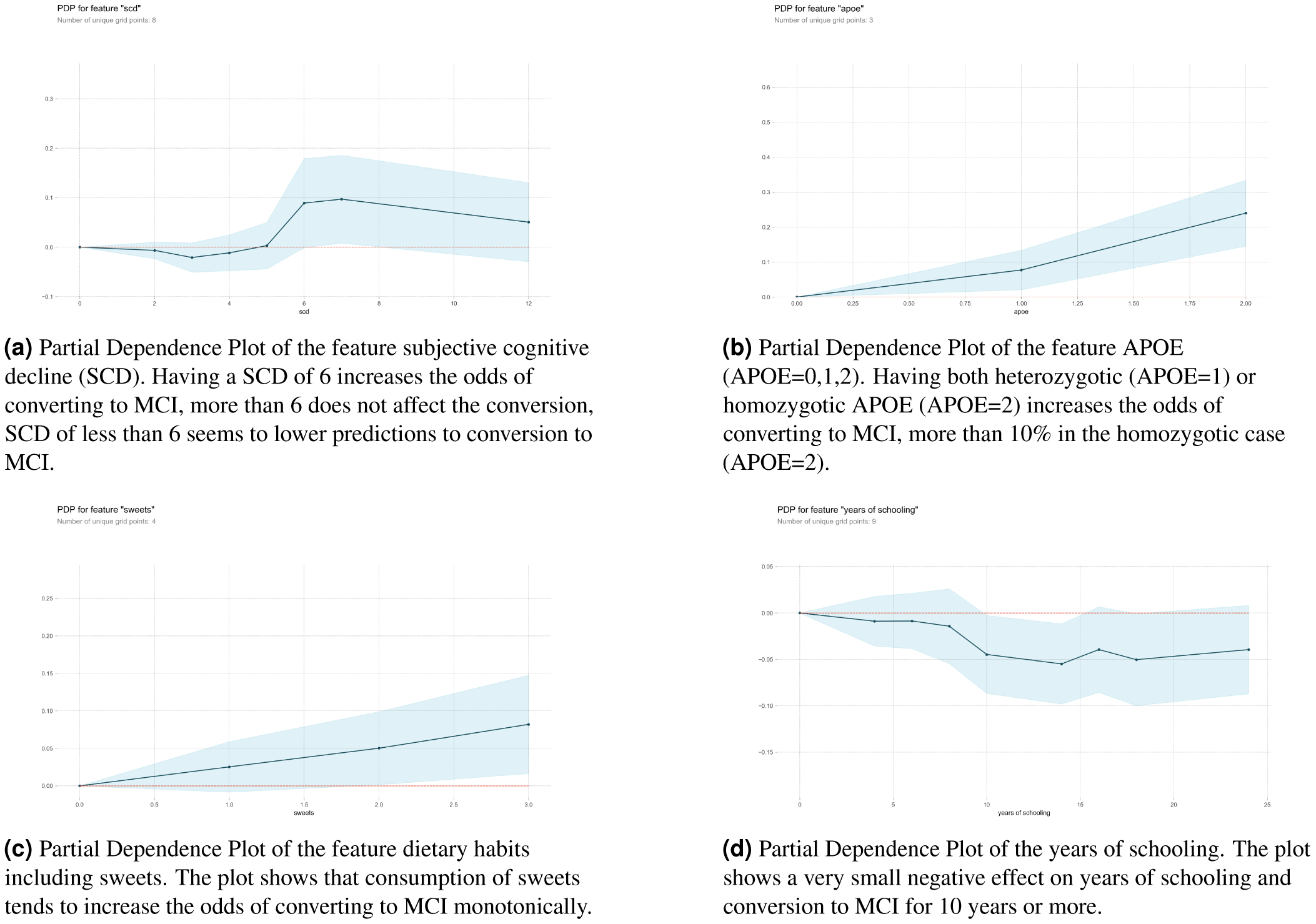
Partial Dependence Plots (PDP) of the variables identified SCD, sweets dietary habit and years of schooling. PDP allow to look inside the range of values of the variable and estimate the effect in the increase/decrease of conversion to MCI for specific values of the feature.

The last permutation-based method borrows from cooperative game theory which is a game between groups or coalitions of players that focuses on predicting the collective payout, this is in contrast with non-cooperative Nash equilibria of individual players^29^. When a machine learning model yields a prediction for an observation, not all the features have equal weight in the prediction, some features may be very important while others are irrelevant. As we have seen above, it is always to estimate the effect of a single feature by removing it or shuffle its values, the bigger the change in the model’s output the larger the role played by the feature in the prediction. However, in this discovery process, the dependencies between features are not being considered. To take into account the interaction among features we cannot single out features but we can use each possible subset of features to study how the prediction changes. In this way, we can determine the unique contribution of each feature without breaking the interdependencies among them.

Shapley values show how much a given feature changed the prediction compared to the prediction at the baseline value of that feature. Shapley (or SHAP for short) values allow us to decompose any particular prediction into the sum of the effect of each feature in that particular prediction. Thus, feature contributions can be positive or negative, a Shapley positive value indicates that the feature acts as a force that pushes the prediction towards “1” (conversion to MCI) while a negative value reflects a force working on the opposite direction, towards “0” (non-conversion to MCI).

Figure 6 plots the SHAP values of every feature for every new sample (184 subjects or 20% in total included in the test set). Figures a) and b) show in the vertical axis the features ranked by importance (top to bottom) calculated as the sum of the SHAP value magnitudes over all samples. The horizontal axis in a) represents the impact on the model prediction (0: no impact, *SHAP* > 0: push towards “1” (conversion to MCI) and *SHAP* < 0 push towards “0” (non-conversion)). The horizontal axis in b) represents the average impact of the SHAP value calculated with the distribution from the left side figure. The figure reveals, for example, that high values of SCD, age, APOE, and sweets increase the prediction of conversion to MCI. The most important feature is again the subjective cognitive decline (SCD).

**Figure 6.**
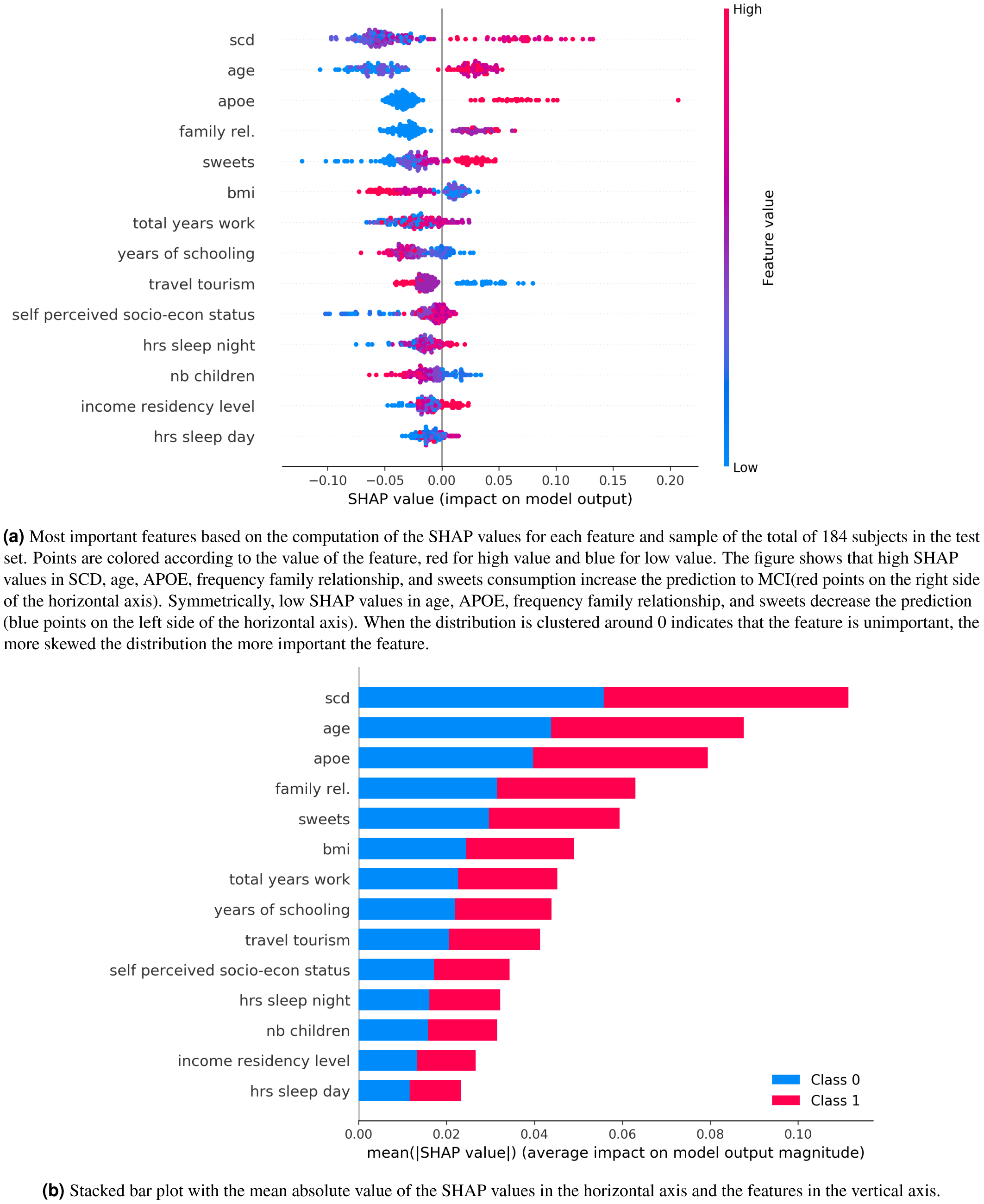
The five most important features are SCD, age, APOE, frequency family relationship family relationship and sweets eating habits. a) Each point represents a subject and b) is the aggregate of the Shapley values.

## Discussion

There is not an iron law of progression from normal to MCI to dementia, some people with MCI might never get worse, and a few would even get eventually better^30^. In this work, we have studied the most important self assessed features for MCI conversion using *The Vallecas Project* Dataset. The rationale and motivation behind solving this feature selection problem is twofold. First, to reduce the complexity in order to gain in interpretability. The *Vallecas Project* dataset has thousands of variables possibly including many redundant and irrelevant features. Second, by identifying lifestyle and other self-assessed features related to MCI conversion, it is possible to provide recommendations and to suggest healthy-aging effective choices.

The feature selection problem of choosing the self-assessed features that contribute most to the target feature (conversion to MCI) was investigated using three different techniques. First, *Filter method* or univariate selection to find the variables that contain the most information about the target variable. Second, *embedded method*, specifically, random forest a learning algorithm with feature selection decision integrated in the learning algorithm. And last, *permutation-based* methods, including random shuffling, partial dependence plot and Shapley values^31^. *Filter methods* pick up the intrinsic properties of the features estimated via the univariate correlation matrix. In *Embedded methods* (random forest) the selection of the feature is integrated with the classifier itself rather than being decided from the external accuracy metric. And finally, *permutation-based* methods assess the importance of features by studying how shuffling the feature values affect the model’s accuracy, important features, when shuffled, will likely make the model predictions to drop.

The strongest result in this study is that the most important self-assessed variable to predict future conversion to MCI is the subjective cognitive decline (SCD). This feature is selected as the most important across all the techniques implemented here for future selection. SCD refers to a self-experienced persistent decline in cognitive abilities in comparison with a previously normal status and independently of the objective performance on neuropsychological tests. This condition has been proved to precede MCI due to AD and thus it is considered that SCD may occur at the final stage of the preclinical phase of AD^32,33^. A recent meta-analysis has shown that almost 25% of older adults who report SCD will develop prodromal AD four years later^34^. In addition to this, the rate of progression to dementia among those individuals who report SCD is twofold during a 5 year following period^35^. All these findings confirm that cognitive complaints perceived by patients themselves are highly informative and should be carefully considered in clinical settings.

This work has important limitations that need to be laid out. When building machine learning or any other statistical inference decision making, the performance of the model prediction can suffer from at least three major factors. First, the predictors do not contain relevant information about the target that needs to be predicted. Second, the model fits poorly the data, for example, it has memorized the data producing overfitting and lastly, the data set is unbalanced i.e. the data do not have enough samples of the minority class.

The feature selection problem studied is affected by all these three issues. The predictors are self-assessed by the participants which logically introduces noise due to the subjective nature in the data collection process. While the accuracy of the random forest is very good especially considering that we are predicting conversion to MCI in a time window from 1 to 5 years, the model performance deteriorated strongly using other metrics like recall or precision. A likely rationale for this circumstance is the unbalanced data set, the ratio of non-converter/converter is 0.82*/*0.12. It is entirely possible that we do not have enough cases of converters for the classifier to extract a consistent pattern. The dataset used here is small 900, leaving a test set of less than 200 samples and far from the sample size in Big Data which go from the order of 10,000 to millions.

Although the problem at hand is particularly challenging, it is however, worth emphasizing that the deliberate choice of looking at the space of self-assessed features brings the possibility to intervene on those factors. For example, dietary and sleeping habits identified as important features for MCI conversion are lifestyle factors that are suitable to be changed. More importantly, we use a set of permutation-based techniques that allow us to estimate the importance of features relative to its range of values and in individual basis, that is to say, we do not only open the black box in identifying which are the most important features in the machine learning algorithm but we also look inside each feature to identify how variations in the feature values affect the prediction, increasing or decreasing the chances to convert to MCI.

A biomarker is a measurable indicator of some biological state. This same idea can be carefully extended to identify lifestyle factors that influence the deterioration of cognitive performance and ultimately the development of AD^36^. The widespread use of smartphones and other smart devices able to collect lifestyle information is expected to foster a more participatory and personalized treatment of chronic diseases. Self-reported data are called to play a key role in the promotion of healthier choices, promoting patients empowerment. SCD measurements are a non-invasive and low-cost method for screening that can be easily included in personal health record systems.

The epistemological implications of the data-driven approach of machine learning are at this point still unclear, however, what is incontestable is that research in problems lacking a clear theoretical framework such as chronic diseases will bring a more intensive (never less) use of data and predictive analytics. Building prediction machines for conversion to MCI using self-informed features, i.e. variables that can be reported by the subject herself, represents a step forward towards a new medicine that aspires to be closer to P4 medicine^37^: *Predictive, Preventive, Personalized* and *Participatory*.

## Acknowledgements

We wish to thank all the participants in *The Vallecas Project* and the staff of Fundación CIEN (Centro de Investigación de Enfermedades Neurológicas).

## Author contributions statement

J.G-R. conceived the study in discussion with MA.F-B. and M.A-V; J.G-R. conducted the analyses; MA.F-B. and M.A-V provided text/references on specific sections. All authors contributed to the text of the manuscript and reviewed the final submission.

## Additional information

### Competing interests

The authors declare no competing interests.

## Supplementary Material

### Self-assessed features

**Table S1.**
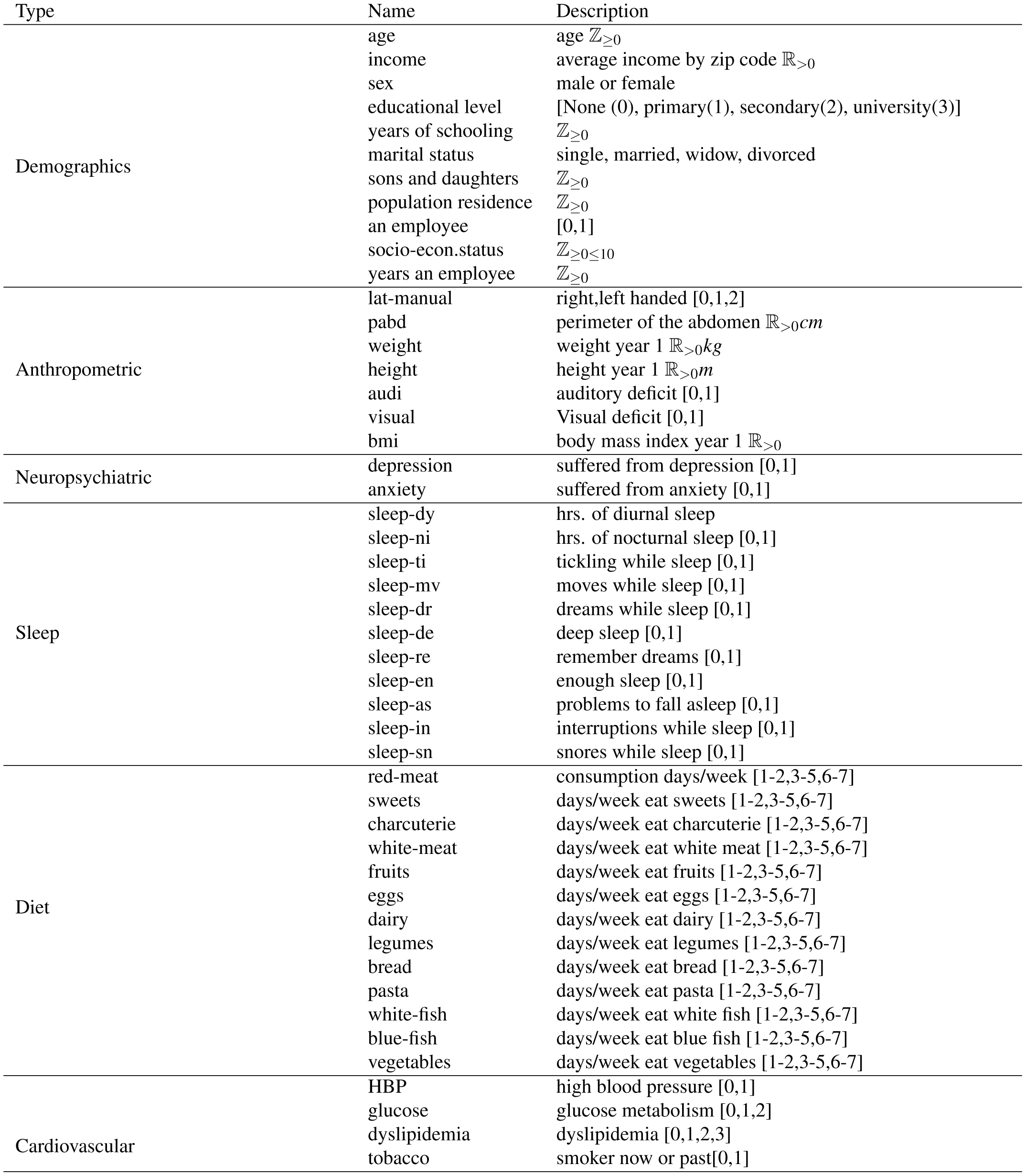
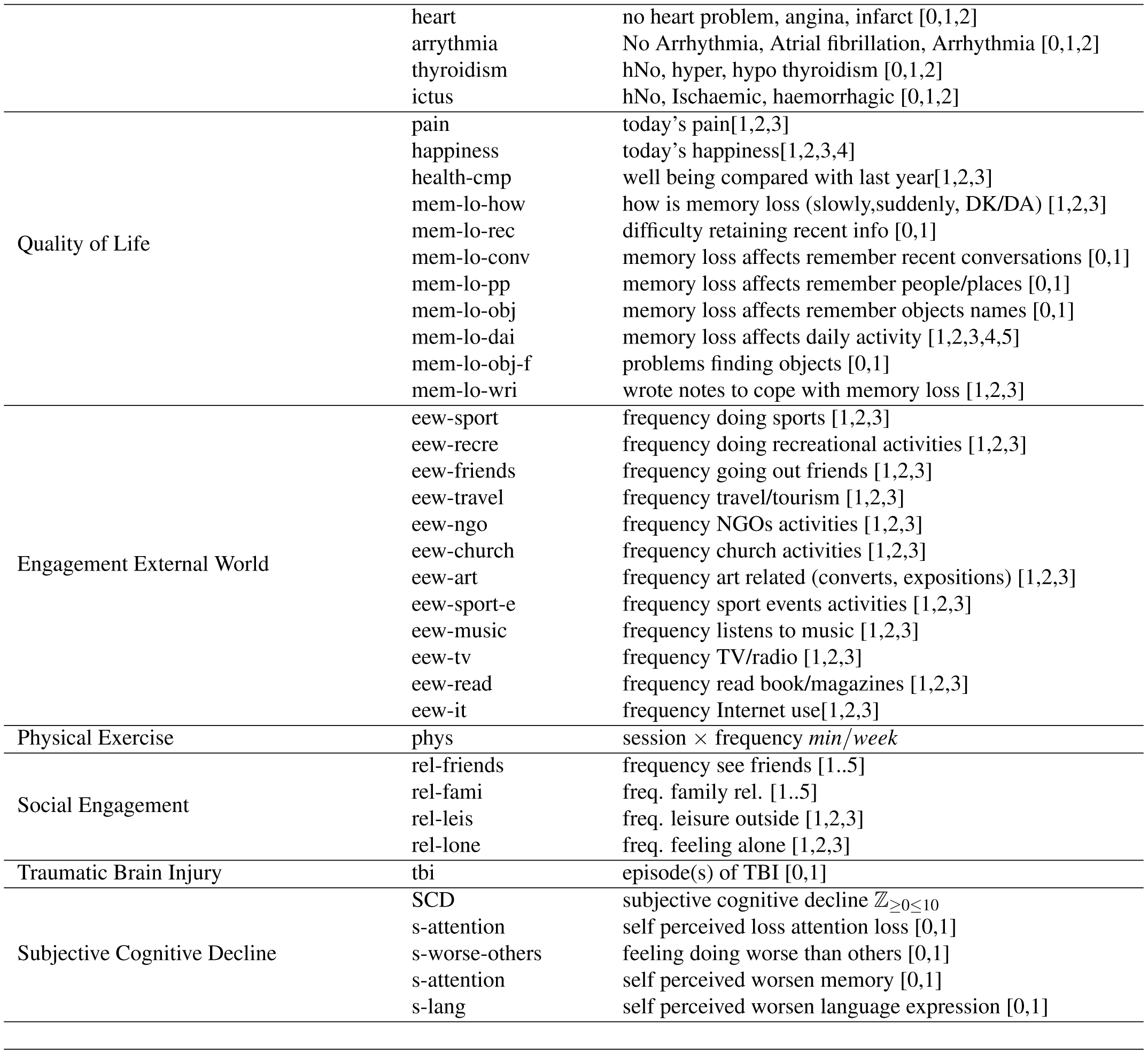
Self-assessed features collected in *The Vallecas Project*

Low-variance features (training-set variance lower than the 20% threshold) are removed. Feature *a13* (use of information technologies IT) is removed since it is strongly correlated with *years of schooling, eqm10* and *eqm83* are also removed since they are correlated with *scd (subjective cognitive decline)*, finally educational level is removed since is strongly correlated with total number of schooling years.

### Random Forest

Whenever we build a random forest we need to tune the hyperparameters which need to be adjusted in order to optimize the desired performance metric. Hyperparameters are outside the model in the sense that are set by the modeler before training. Note the difference with model parameters which are learned during training. The hyperparameter tuning consists in K-Fold (*K* = 5) cross validation, that is, we split the training set into K folds (subsets of the training set), then we iteratively fit the model *K* times, each time training the data on *K* − 1 folds and evaluating on the *K*-th fold. To find the best hyperparameters we use a dual approach, first we use randomized search to randomly sample from the grid of hyperparameter range. The set of hyperparameters returned in the randomized search is used to inform the Grid search method run afterwards and that exhaustively searches all possible combination of hyperparameters.

The set of optimal hyperparameters obtained are shown in Table S3.

**Table S3.** Hyperparameters of the Random Forest Classifier using Grid Search cross validation. (Suplementary Materials Figure S1 shows one tree out of the 100 trees built in the forest).

| Hyperparameter | Value | Description |
| --- | --- | --- |
| bootstrap | True | Whether bootstrap samples are used when building trees. If False, the whole dataset is used to build each tree. |
| class weight | balanced | weight associated with each class |
| criterion | Gini | function to measure the quality of split |
| max depth | 3 | maximum depth of the tree |
| max features | 2 | number of features to consider for the best split |
| min samples leaf | 0 | minimum number of samples required to be at a leaf node |
| min samples split | 2 | minimum number of samples required to split an internal node |
| estimators | 100 | total number of trees in the forest |

Inherent in the hyperparameter tuning process is the evaluation criterion used. The evaluation criterion consists in computing scoring objects that gives us information about model performance, for example, accuracy.

For the sake of illustration, Figure S1 shows one tree out of the 100 trees built in the forest. The important features in a decision tree are located in the nodes close to the root of the tree and the unimportant ones will tend to be close to the leaves of the tree or entirely absent from the tree. Therefore, random forests allow us to get an estimate of the importance of any feature by calculating how deep in the tree the feature appears across all the trees. Specifically, feature importance is calculated as the decrease in node impurity weighted by the probability of reaching that node. The mean decrease in impurity importance of a feature is computed by measuring how effective the feature is at reducing uncertainty (classifiers) or variance (regressors) when creating decision trees within Random Forest.

**Figure S1.**
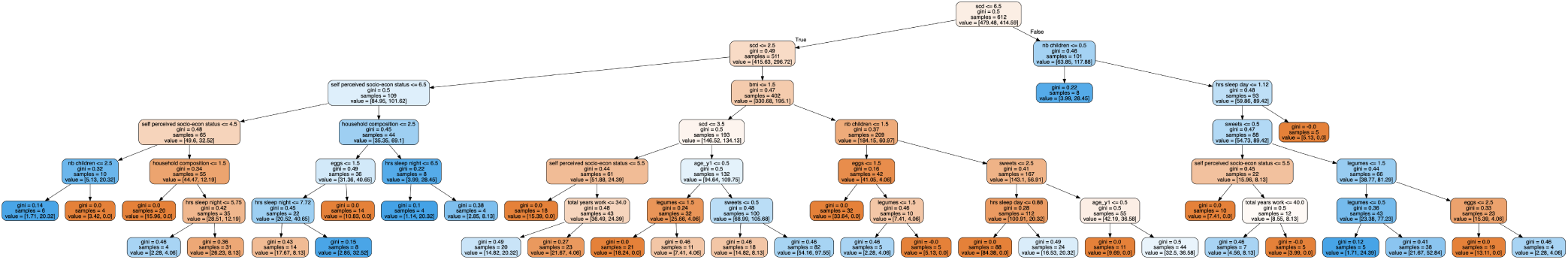
The figure shows one tree of the random forest. The root of the tree is the node with the highest Gini score, subjective cognitive decline (SCD). The nodes closer to the root are more important than those at the bottom of the tree as the Gini value included in each mode indicates. The maximum depth of the tree is 6. Nodes with red color refer to samples that fall into the group of non converter to MCI, the boxes with blue color groups the converters.

### Shapley Value

The idea behind the Shapley value is that each feature value is a player in a prediction game and the game’s payout is the accuracy of the prediction. For example, the prediction *C* for two features *X* ={*X*_1_, *X*_2_} according to the function *f*(*X*) = *C* is described in the bellow table. We want to compute the contributions of each feature for a given observation e.g. y=(0,1).

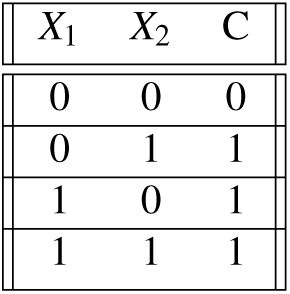

| $X_1$ | $X_2$ | C |
| --- | --- | --- |
| 0 | 0 | 0 |
| 0 | 1 | 1 |
| 1 | 0 | 1 |
| 1 | 1 | 1 |

First, we need to compute the expected prediction if no feature values are known, and from that we can compute what we need which is the prediction differences for all subsets of features {∅},{1},{2},{1, 2}}

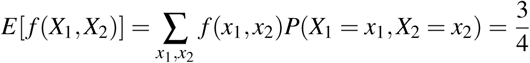

The prediction differences are then:

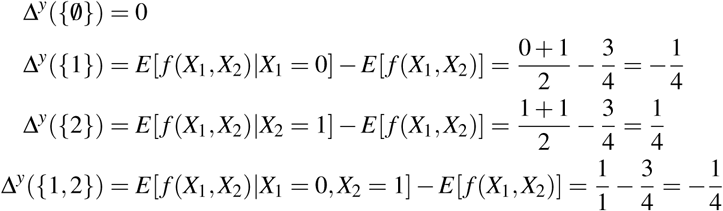

The last step is to calculate the contribution of each feature *X*_2_ and *X*_2_ using the formula of Shapley value shown in 3

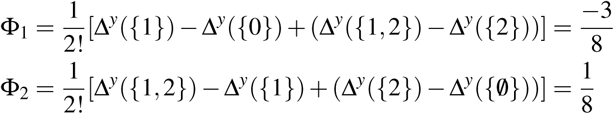

Feature *X*_2_ has a positive influence because it made the model predict 1, feature *X*_1_, on the other hand has a negative contribution because it made less probable to predict 1. Also, feature *X*_1_ is larger in absolute value and therefore is more important for the prediction than *X*_2_. To summarize, the Shapley values Φ_1_ and Φ_2_ tells us that the model was influenced by both features for the prediction of the instance (0,1), with *X*_1_ being more important than *X*_2_, being *X*_1_ against and *X*_2_ in favor of the decision.

